# Why should mitochondria define species?

**DOI:** 10.1101/276717

**Authors:** M.Y. Stoeckle, D.S. Thaler

## Abstract

More than a decade of DNA barcoding encompassing about five million specimens covering 100,000 animal species supports the generalization that mitochondrial DNA clusters largely overlap with species as defined by domain experts. Most barcode clustering reflects synonymous substitutions. What evolutionary mechanisms account for synonymous clusters being largely coincident with species? The answer depends on whether variants are phenotypically neutral. To the degree that variants are selectable, purifying selection limits variation within species and neighboring species may have distinct adaptive peaks. Phenotypically neutral variants are only subject to demographic processes—drift, lineage sorting, genetic hitchhiking, and bottlenecks. The evolution of modern humans has been studied from several disciplines with detail unique among animal species. Mitochondrial barcodes provide a commensurable way to compare modern humans to other animal species. Barcode variation in the modern human population is quantitatively similar to that within other animal species. Several convergent lines of evidence show that mitochondrial diversity in modern humans follows from sequence uniformity followed by the accumulation of largely neutral diversity during a population expansion that began approximately 100,000 years ago. A straightforward hypothesis is that the extant populations of almost all animal species have arrived at a similar result consequent to a similar process of expansion from mitochondrial uniformity within the last one to several hundred thousand years.

## Precis

1. Mitochondrial Cytochrome Oxidase Subunit I DNA barcodes (COI barcodes, often shortened to “DNA barcodes” or “barcodes” in this article) began as an aid to animal species identification and made no claims of contributing to evolutionary theory. Five million DNA barcodes later the consistent and commensurable pattern they present throughout the animal kingdom is one of the most general in biology. In well-studied groups the majority of DNA barcode clusters agree with domain experts’ judgment of distinct species.
2. The tight clustering of barcodes within species and unfilled sequence space among them are key facts of animal life that evolutionary theory must explain. Many aspects of speciation are complex. Barcodes are unique in being quantifiably commensurable across all animal species and almost always yielding the same single simple answer [1].
3. Either of two evolutionary mechanisms might account for the facts: a) species-specific selection, or b) demographic processes acting independently of phenotype.
4. Most barcode variation consists of synonymous codon changes. Since the assumption of neutrality of mitochondrial synonymous codons was asserted, many exceptions in nuclear genes and prokaryotic systems have been found.
5. New arguments are presented that synonymous codon changes in mitochondrial genes are neutral to a greater extent than nuclear genes.
6. Extensive data on modern humans make our species a valuable model system for animal evolution as a whole. The mitochondrial variation within the modern human population is about average when compared to the extant populations of most animal species.
7. Similar neutral variation of humans and other animals implies that the extant populations of most animal species have, like modern humans, recently passed through mitochondrial uniformity.

## History of COI barcoding

DNA barcoding was first proposed as a tool for practical taxonomy and to democratize actionable biological knowledge [2, 3]. At its origin DNA barcoding made no claim of contributing to evolutionary theory. Previous work bode well for mitochondrial genomes being reliably similar within animal species yet in many cases distinct among neighbor species [4, 5]. The particular mitochondrial sequence that has become the most widely used, the 648 base pair (bp) segment of the gene encoding mitochondrial cytochrome c oxidase subunit I (COI), reached a tipping point because widely applicable reliable primers and methods useful for both vertebrates and invertebrates were adopted by a critical mass of the community [6, 7].

Skeptics of COI barcoding [8] raised a number of objections about its power and/or generality as a single simple metric applicable to the entire animal kingdom, including: 1) the small fraction of the genome (about 5% of the mitochondrial genome and less than one millionth of the total organism’s genome) might not be sensitive or representative [9, 10]; 2) since animal mitochondria are inherited maternally the apparent pattern of speciation from mitochondria is vulnerable to distortion when females and males roam differently [11]; 3) the mitochondrial chromosome is subject to types of selection not experienced by the nuclear genome [12]: replicon competition within each organelle [13], among organelles inside each cell [14–16], including differential segregation of organelles at cell division [17]; and 4) mitochondria in some groups are sensitive to agents such as *Wolbachia* that are not known to affect nuclear genes [18]. Mitochondrial pseudogenes in the nucleus sometimes confused analysis [19]. Anecdotally, some domain experts felt that only specialists can reliably recognize species in each group and that “DNA taxonomy” was felt as necessarily inferior or a threat.

The current field of COI barcodes is no longer fragile but neither is it complete. As of late 2016 there were close to five million COI barcodes between the GenBank and BOLD databases. Objections can now be seen in the cumulative light of these data and more than a decade’s experience. There is no longer any doubt that DNA barcodes are useful and practical (Figs. 1,2). The agreement with specialists encompasses most cases in several important animal domains. Many cases where DNA barcodes and domain specialists do not agree reflect geographic splits within species or hybridization between species. Others upon further investigation been attributed to mislabeling or sequence error [20]. Some may represent *bona fide* exceptions to the rule that mitochondrial sequence clusters coincide with species defined by other means. In the great majority of cases COI barcodes yield a close approximation of what specialists come up with after a lot of study. Birds are one of the best characterized of all animal groups and COI barcode clusters have been tabulated as agreeing with expert taxonomy for 94% of species [21].

**Fig. 1.**
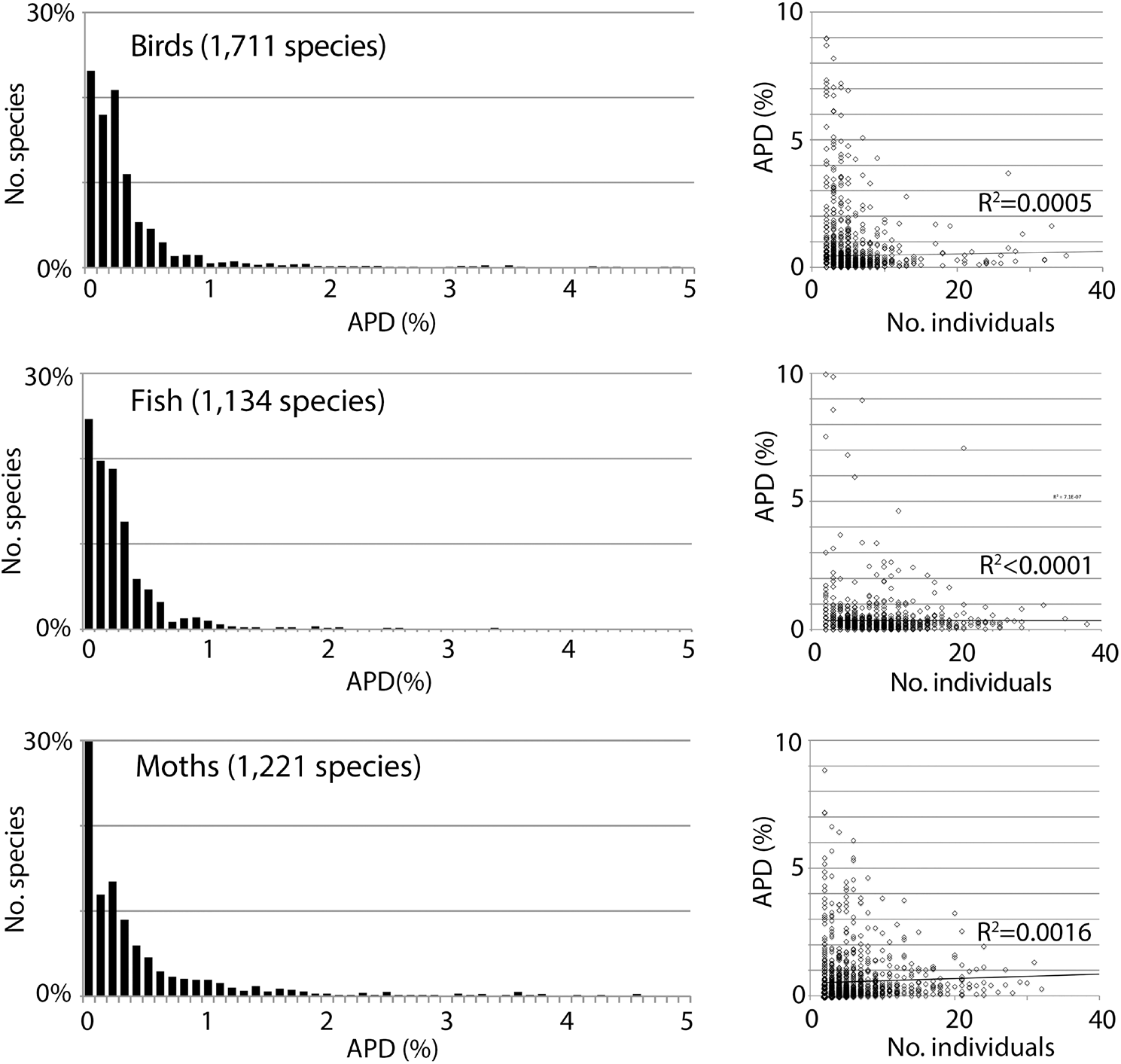
**Low intraspecific COI barcode variation is the norm in animals**, not an artifact of handpicking examples or small sample size. Variation is expressed as average pairwise difference (APD) between individuals.

**Fig. 2.**
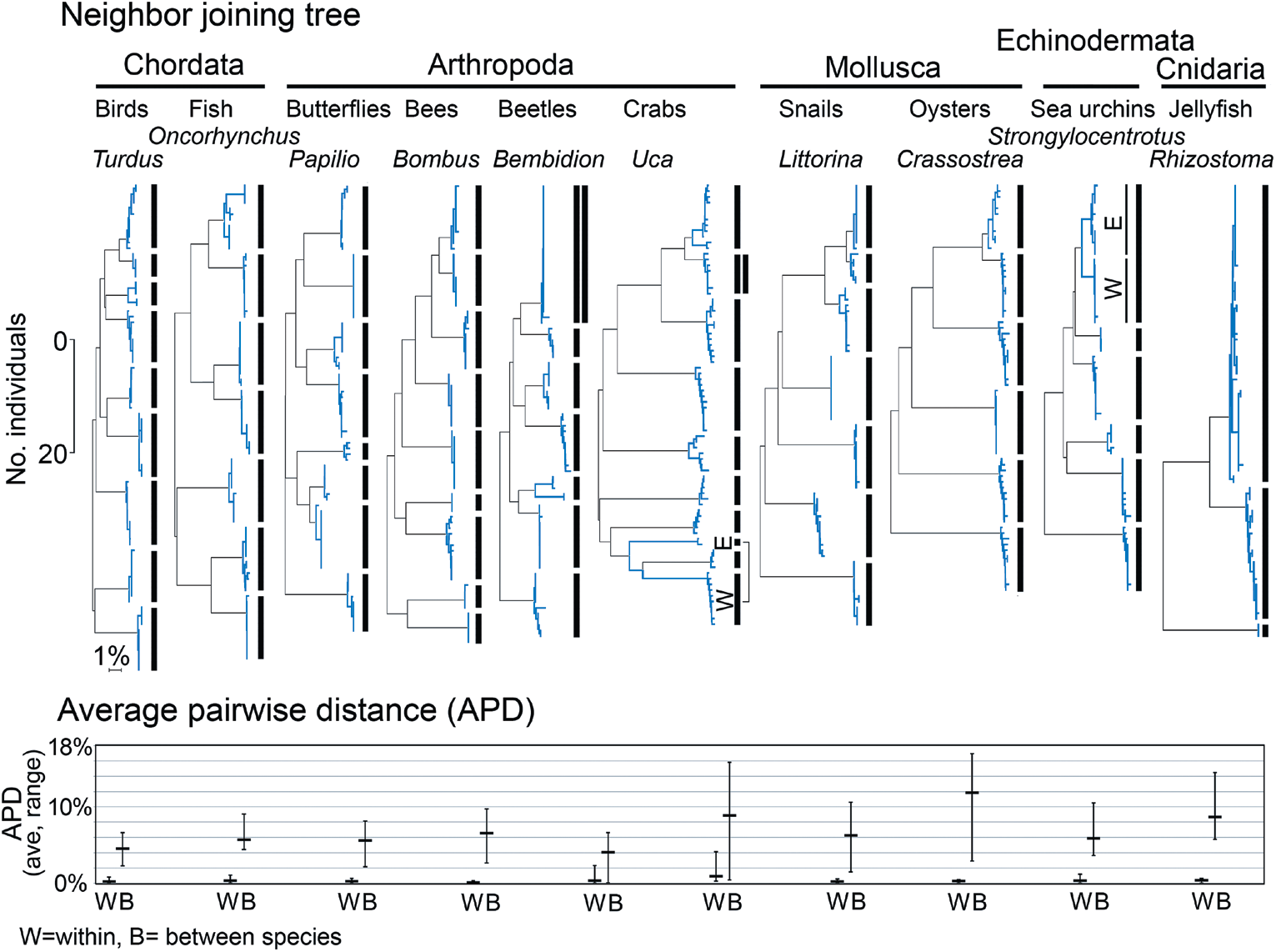
Relatively large interspecific differences, as compared to uniformly small intraspecific differences, are the norm in animals. Together these yield the familiar clustering pattern that enables DNA barcode species identification. Shown are neighbor-joining (NJ) trees (with scale bars for number of individuals and percent K2P distance) and average pairwise distance (APD) within and between sets of closely-related congeneric species. At top, NJ trees with bars marking species clusters. Exceptions to the one species/one cluster rule include cases with multiple clusters within species, corresponding to geographically isolated populations [marked as (W)estern and (E)astern], and cases with clusters shared between species, marked by double vertical lines. At bottom, APDs for the same congeneric sets, with average (horizontal bar) and range (vertical bar) of intraspecific and interspecific APDs shown.

## Exceptions to the rule that each species is a single unique cluster

Most exceptions to the generality that COI clusters represent species are also exceptions to the general rule that species are single interbreeding populations. These include cases with phylogeographic divisions within species and those with shared or overlapping barcode clusters (Figs. 2,3).

**Fig. 3.**
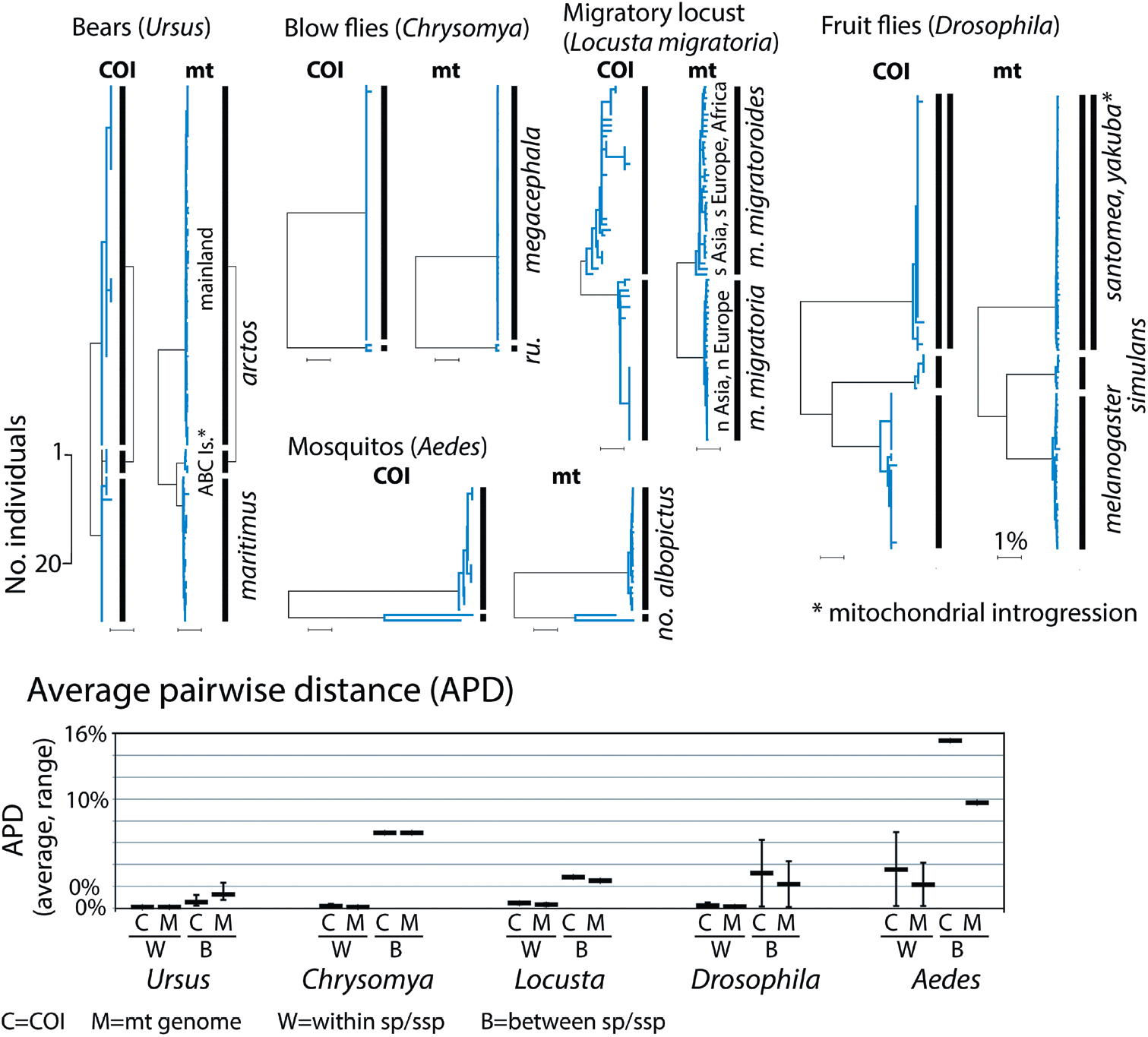
Clustering of 0.6 kbp COI barcode segments accurately represents the complete 12 kbp coding mitogenome. At top, COI and mt genome NJ trees exhibit similar clustering patterns. At bottom, average pairwise differences within and between species in each set are about the same whether calculated from COI barcodes or coding mitogenomes. As in Fig. 2 legend, apparent exceptions with phylogeographic divisions (locusts) or shared or overlapping clusters (bears, fruit flies) are noted. NJ tree scale bars for number of individuals and percent K2P distance are shown.

In most well-studied cases of shared or overlapping barcodes, nuclear genome analysis demonstrates these anomalies are due to hybridization resulting in mitochondrial introgression from one species into the other. If recent, and complete across the whole population, introgression erases mitochondrial differences between species. Introgression events in the more distant past and those involving only part of a species produce more complex patterns, as illustrated by *Ursus* bears (Fig. 3). Based on nuclear and mitochondrial genome analysis, polar bears *(U. maritimus)* hybridized with “ABC island” brown bears *(U. arctos)* about 50,000 years ago, with introgressive replacement of ABC *arctos* mitogenomes by *maritimus* mitogenomes. The mitochondrial lineages subsequently diverged, but ABC island brown bear mtDNA remains closer to polar bear than to mainland brown bears. Nuclear genomic analysis supports taxonomic classification of ABC island and mainland populations as subspecies of brown bear.

Incomplete lineage sorting with retention of ancestral polymorphisms is a plausible mechanism for shared or overlapping mitogenomes, also called paraphyly. However, in all cases we know of, when analyzed for nuclear and mitochondrial differences, ongoing or historical hybridization is the likely cause (see Fig. 1 and Table S3 in reference [20]).

## General across the animal kingdom

DNA barcodes based on mitochondrial sequences might have failed to be sensitive, general, practical, or to agree with the judgment of experts in each domain. Five million DNA barcodes later some exceptions have been found, however, the power, generality, and validity of the COI barcode approach for identifying animal species is no longer in question, at minimum, for several major groups (Figs. 1–3). A general observation is that barcode clusters correspond best to species in well-studied animal groups, where taxonomists have mostly decided and agreed upon what species are. Thus there is good support in several major phyla, including Chordata, Arthropoda, Mollusca, Echinodermata. We note that these phyla are estimated to contain about ¾ of named animal species.

## Incompletely studied groups

In the remaining 23 animal phyla, there are examples where clusters match species, but the overall picture is muddier. Many are small animals, difficult to distinguish morphologically, and have attracted relatively little taxonomic or DNA barcode study. Major incompletely studied groups include Annelida, Nematoda, Platyhelminthes, Porifera, and Rotifera. We expect that with further study these phyla will fit a pattern similar to that in more established groups. However, at this stage it takes cherry-picking to find examples that match the better-studied phyla and one cannot make a data-based case for the general validity of DNA barcoding in these phyla.

Beyond using the DNA barcode as an aid to taxonomy, the enormity of data now available make it appropriate to extend the applications of the “broad but not deep” vista that COI barcodes uniquely provide [20, 22, 23]. In a founding document of phylogeography, Avise and colleagues noted the long-standing divide in biology between the intellectual lineages of Linnaeus for whom species are discrete entities and those of Darwin who emphasize incremental change within species leading to new species [4]. They presciently proposed that mitochondrial analysis would provide a way to bridge the intellectual gap. DNA barcoding now provides the most comprehensive database allowing a kingdom-wide and quantitative realization of that vision.

## Differing definitions of species

There are approximately 30 different definitions of species in the biological, philosophical, and taxonomical literatures [24]. Almost all of them share the idea that species are distinct entities in biology and the corollary idea that there are discontinuities among species [25]. In their clarifying and valuable analyses Mayr [25] and de Queiroz [11] point out that all definitions of species involve separate monophyletic evolutionary lineages (with important exceptions where symbiosis or horizontal gene transfer are key [26]). Different distinguishing factors such as mating incompatibility, ecological specialization, and morphological distinctiveness evolve, in various cases, in a different temporal sequence. During the process, as species diverge and emerge some of these characteristics will be fulfilled while others are not. Disagreement is inevitable when different properties are considered necessary and sufficient to fit one or another definition of “species”.

There are two important observations regarding how COI barcodes fit into the differing definitions of species. First, the cluster structure of the animal world found in COI barcode analysis is independent of any definition(s) of species. Second, domain experts’ judgments of species tend to agree with barcode clusters and many apparent deviations turn out to be “exceptions that prove the rule”. Controversy around the edges, e.g. disagreements about whether or not borderline cases constitute species or subspecies [27, 28] should not obscure visualizing the overall structure of animal biodiversity. It is unavoidable that some cases will be considered as species by one definition and not another. Controversial cases can illuminate in the context of William Bateson’s adage to “treasure your exceptions” [29] but they should not obscure the agreement for most cases and an appreciation of the overall structure within the animal kingdom. This pattern of life, close clustering within individual species with spaces around clusters, can be visualized and demonstrated in different ways and with different statistics (e.g., Figs. 1-4). It qualifies as an empirically-determined evolutionary law [30]. Barcode distribution is arrived at independently but consistent with a view of biology as composed of discrete entities that on different levels include organisms [31] and species [32].

**Fig. 4.**
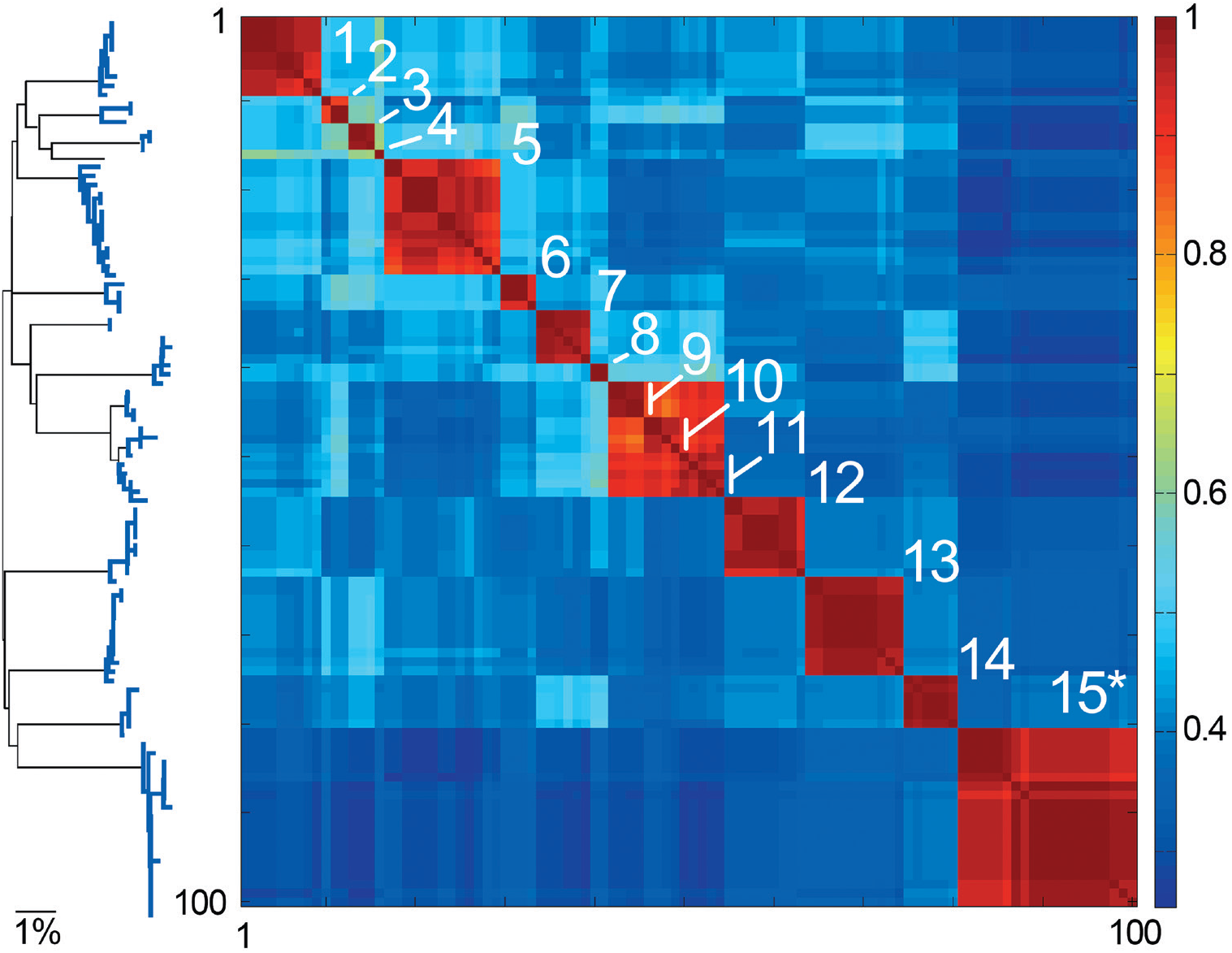
Species are islands in sequence space. COI barcode NJ tree and Klee diagram of American Robin *(Turdus migratorius)* and closely related *Turdus* species. To generate dataset, a single American robin COI barcode was used to search GenBank using BLAST, and the top 100 matches were downloaded. In Klee diagram, numbers indicate species, asterisk marks *T. migratorius* sequences, and indicator vector correlation scale is at right, with 1 representing 100% sequence identity.

## The pattern of DNA barcode variance is the central fact of animal life that needs to be explained by evolutionary theory

In ‘The Structure of Scientific Revolutions’ Thomas Kuhn makes the point that every scientific model takes certain facts of nature or experimental results as the key ones it has to explain [33]. We take the clustering structure of COI barcodes—small variance within species and often but not always sequence gaps among nearest neighbor species—as the primary fact that a model of evolution and speciation must explain. The pattern of life seen in barcodes is a commensurable whole made from thousands of individual studies that together yield a generalization. The clustering of barcodes has two equally important features: 1) the variance within clusters is low, and 2) the sequence gap among clusters is empty, i.e., intermediates are not found. Beyond the qualitative descriptor “low” for the variance within species there is a quantitative statement. The average pairwise difference among individuals (APD; equivalent to population genetics parameter π) within animal species is between 0.0% and 0.5%. The most data are available for modern humans, who have an APD of 0.1% calculated in the same way as for other animals (See Fig. 2 in [34] and Fig. 7 in this paper).

**Fig. 5.**
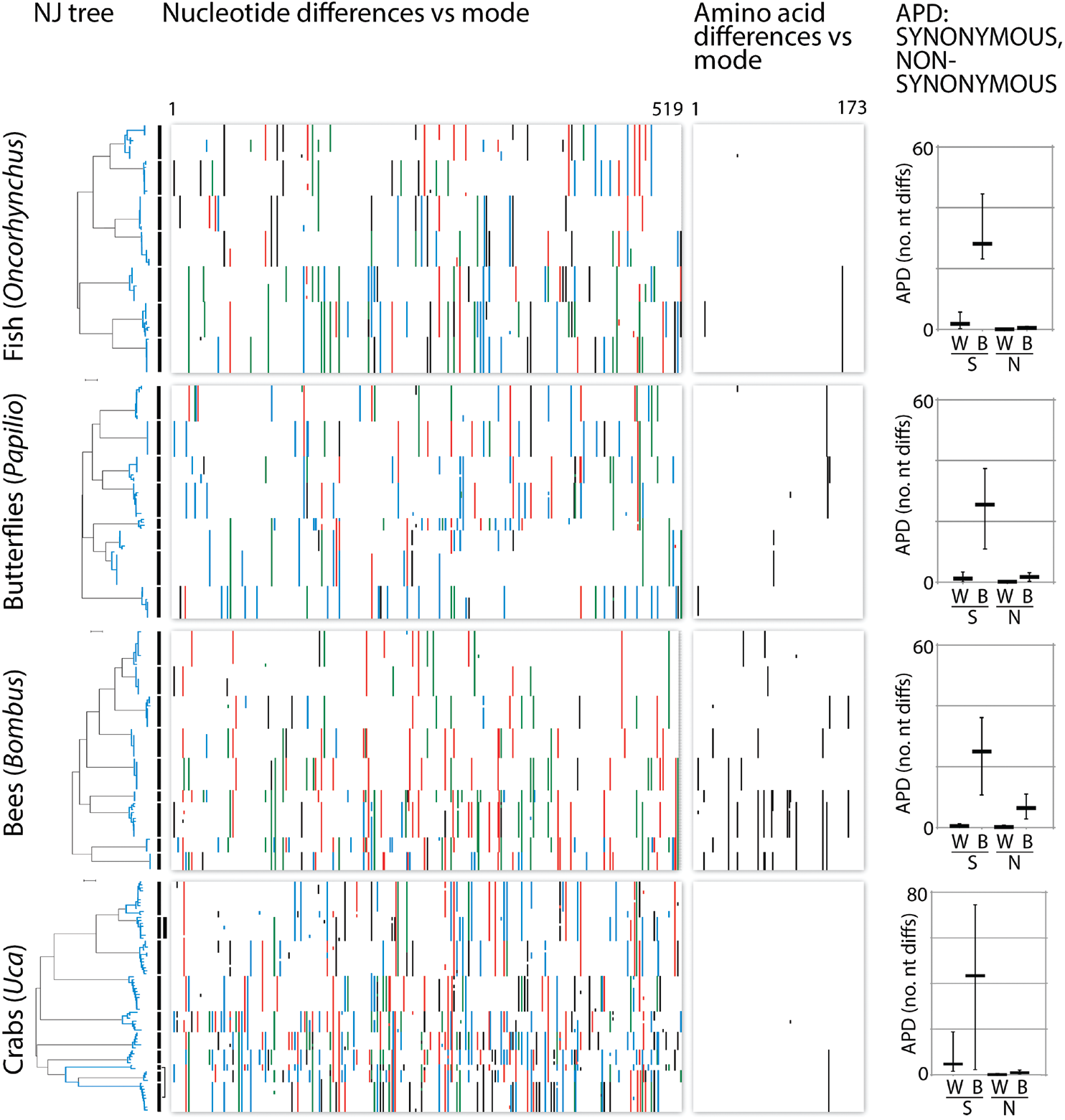
mtDNA clusters reflect synonymous substitutions. Charts depict nucleotide and amino acid differences from the mode for congeneric COI barcode sets in Fig. 3. Nucleotide differences are colorized (A=green; C=blue; G=black; T=red). To minimize contribution of sequence errors and missing data, the 648 bp barcode region is trimmed by 10% at either end, leaving 519 nt/173 amino acids. At right, synonymous (S) and non-synonymous (N) average pairwise distances within (W) and between (B) species. Horizontal bar indicates mean and vertical line indicates maximum and minimum.

**Fig. 6.**
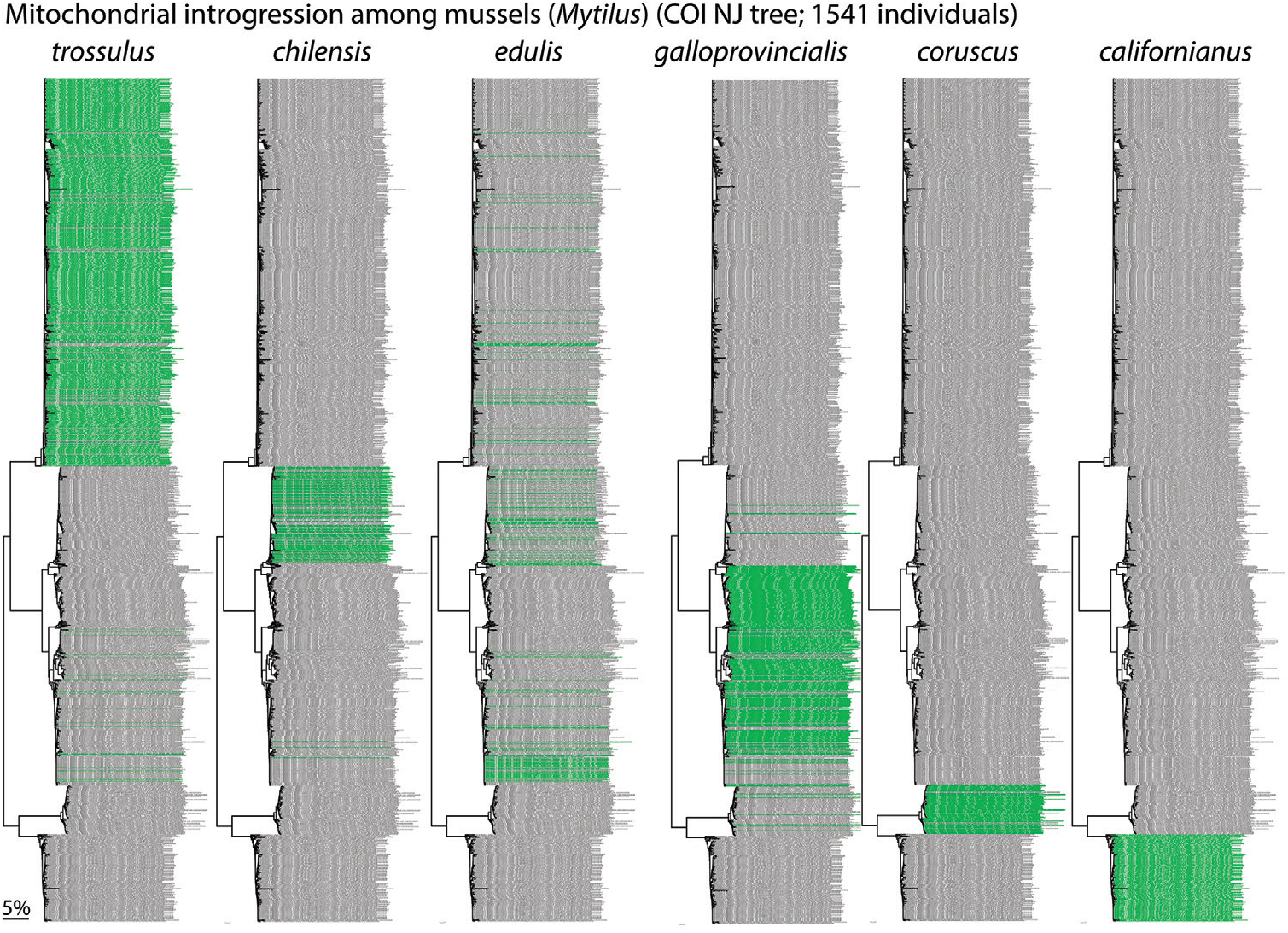
Fertile hybrids. Mytilus mussels exhibit complex patterns of mitochondrial and nuclear introgression, reflecting multiple historical and recent hybridization events, some following introduction of non-native species for aquaculture. F1 hybrids are fertile even though parental species differ by 10-20% in COI nucleotide sequence. This supports view that mtDNA clustering is not due to species-specific adaptations.

**Fig. 7.**
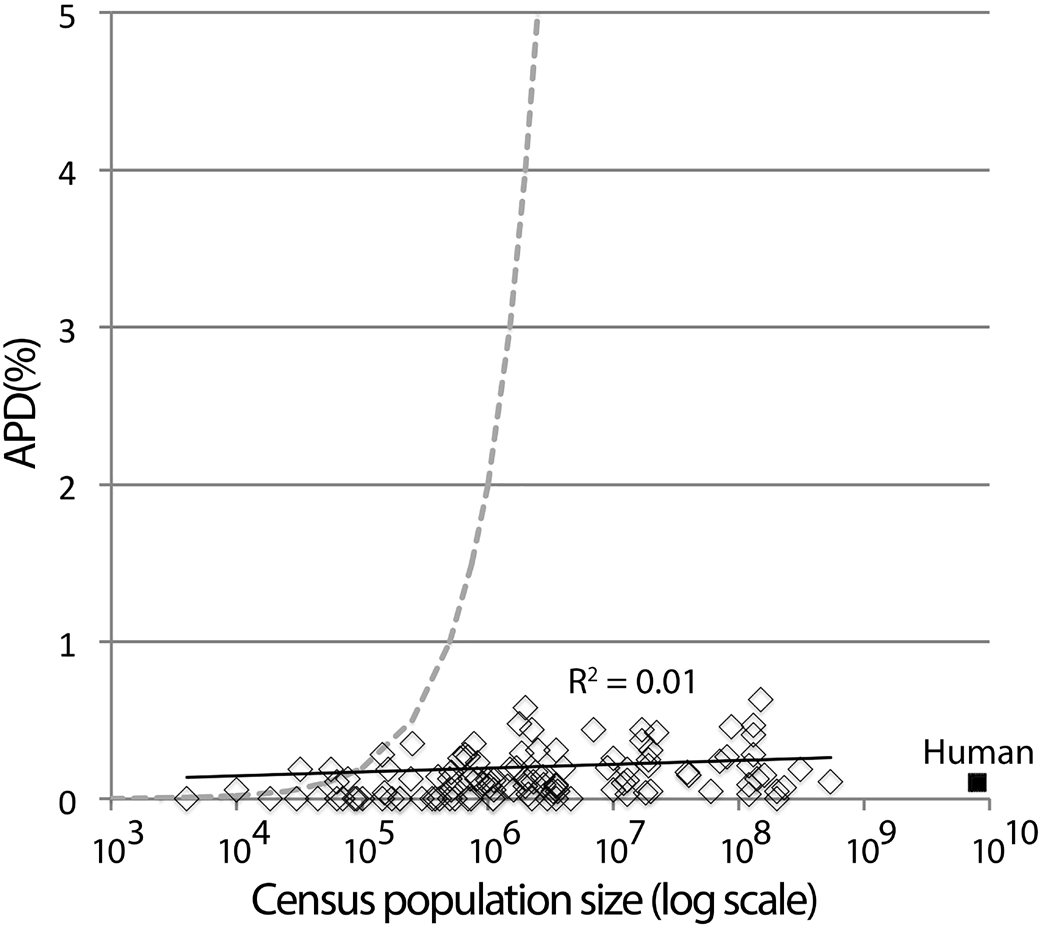
Kimura’s equilibrium model alone is insufficient to account for usual levels of intraspecific variation in animal species. APD and census population size for 112 bird species without phylogeographic clusters are shown. Dashed line is expected APD limit due to (AVP = 2 N μ, where N = population size and μ = mutation rate, using 10^−8^ substitutions/site/ generation, or 1% per My, assuming generation time is 1 y). Average effective population size in the birds shown is 70 thousand (range 0-300 thousand); average census population size is 30 million (range 5 thousand to 500 million). Human mitochondrial variation (population 7.5 billion, APD 0.1%) is typical of that in other animal species.

The agreement of barcodes and domain experts implies that explaining the origin of the pattern of DNA barcodes would be in large part explaining the origin of species. Understanding the mechanism by which the near-universal pattern of DNA barcodes comes about would be tantamount to understanding the mechanism of speciation.

The clustering pattern of life was elegantly articulated by Dobzhansky in his 1937 book Genetics and the Origin of Species [35] from which an extensive quote is merited. Only through DNA barcodes can the same metric be used so that the “feeling that it must be right” can now be given a single quantitative meaning across the entire animal kingdom:

*If we assemble as many individuals living at a given time as we can, we notice that the observed variation does not form a single probability distribution or any other kind of continuous distribution. Instead, a multitude of separate, discrete, distributions are found. In other words, the living world is not a single array of individuals in which any two variants are connected by unbroken series of intergrades, but an array of more or less distinctly separate arrays, intermediates between which are absent or at least rare. Each array is a cluster of individuals, usually possessing some common characteristics and gravitating to a definite modal point in their variation…. Therefore the biological classification is simultaneously a man-made system of pigeonholes devisedfor the pragmatic purpose of recording observations in a convenient manner and an acknowledgement of the fact of organic discontinuity.*

## Two models have the potential to explain the structure of COI barcodes in the extant animal kingdom

Either 1) COI barcode clusters represent species-specific adaptations, OR 2) extant populations have recently passed through diversity-reducing regimes whose consequences for sequence diversity are indistinguishable from clonal bottlenecks. This way of framing the problem is similar to that raised by the analysis of isozymes by electro-phoresis more than half a century previous [36]. The key difference being that the COI barcode data are vast and commensurable across the animal kingdom. “Commensurable” means using the same measurement and being directly comparable. “The “awesome power of’ [37] mitochondrial COI barcodes allows the same metric to be used animal kingdom-wide. Without DNA barcodes, generalizations to all animals have to be based on putting together data sets gathered and analyzed by different methods.

Most barcode variation among neighboring species—and also within species—consists of synonymous codon changes (Fig. 5). The question that determines which of the two mechanisms is most plausible is whether or not synonymous codons in the mitochondrial genome are selectively neutral. If purifying selection does not act on synonymous codons in the mitochondrial genome then demographic processes must be acting to suppress neutral variance.

## Are synonymous codons in mitochondria neutral?

Comparative rates along phylogenies have been used to argue that amino acid changes in the mitochondrial genome are subject to purifying selection but synonymous substitutions are not [23, 38, 39]. Across the animal kingdom the preponderance of SNP variation in mitochondrial sequences consists of synonymous codon changes. Are these synonymous codons targets for purifying and/or adaptive selection strong enough to be responsible for the low variance within species and/or the different consensus sequence among neighboring species? Codon bias in the mitochondrial genome has been shown at the phylogenetic level of order but there is no evidence for different codon bias among neighboring species [40, 41]. Furthermore, the number of synonymous codons relevant to the discussion of DNA barcodes (0.0%-0.5% within species, 0.0%-5.0% for neighboring species) is not enough to alter codon bias. Nearby species do not differ in overall codon bias or GC ratios [40, 41], an observation in contrast to a prediction of the hypothesis that GC bias is an important factor in speciation [42]. If synonymous codons are differentially selected in DNA barcodes, this selection must be acting at the level of the placement of individual codons rather than their cumulative average effects on base composition.

## An assumption of neutrality for synonymous codons is no longer a “slam dunk” [43], i.e., is not a certain conclusion

Avise et al in 1987 made an absolutist statement on the irrelevance of synonymous codons to selective fitness:

*First, in a mechanistic sense, we already “know” that most of the particular mtDNA genotypic variants segregating in populations probably have, by themselves, absolutely no differential effect on organismal fitness. These include, for example, base substitutions in silent positions of protein-coding genes, and some substitutions and small addition/ deletions in the nontranscribed D-Ioop region. These changes are disproportionately common in mtDNA [25] and are ones for which only the most ardent selectionist would argue a direct link to organismal fitness.*

This statement merits critical evaluation in light of three intervening decades of molecular genetics. (Spoiler alert: in this case re-examination strengthens the original assertion for reasons that the authors themselves could not have anticipated.)

Since 1987 numerous examples have emerged where even very few synonymous codon changes make important and selectable differences in organismal, cellular, or viral physiology [44–50].

Synonymous codons may also modulate protein folding or membrane insertion concomitant with translation [51], as suggested for synonymous codons that modulate phenotypes of the cystic fibrosis transmembrane conductance regulator [52] or the p-glycoprotein (multidrug resistance protein) [53]. The cited cases involve cytoplasmic translation of bacteria, bacteriophage, and drosophila nuclear genes. However, they require one to critically re-examine the assertion of absolute and universal neutrality with regard to mitochondria.

## New justification for an old assertion

The proven cases where synonymous codons have phenotypes are all consequent to their differential rate of translation. Codon-specific translation rates are in turn attributed to different concentrations of codon-specific tRNAs, known as “isoacceptors.” This has been proven directly in an experimental system where overexpression of the cognate tRNA changes a low-expression codon to a high one [54, 55]. Isoacceptor concentrations differ between species and even within individuals in a tissue-specific manner [56, 57]. The frequencies of synonymous codons and the concentration of their isoacceptor tRNAs with complementary anticodons coevolve [58]. Codon-specific isoacceptor tRNAs are tightly and dynamically regulated; they play important roles in the differential regulation of gene expression [59]. The human nuclear genome encodes tRNAs with 51 distinct anticodons for the 20 amino acids [60]. In addition to isoacceptors there are dozens to hundreds of “isodecoder” genes in the nuclear genome. Isodecoders are tRNAs that share an anticodon sequence but differ elsewhere. The different sequences of isodecoders are often also associated with different post-transcriptional modifications [60]. It is likely that isodecoders add a further important layer to differential gene expression depending on a codon’s sequence and tissue context but this remains to be proven.

In striking contrast to the multitude of different nuclear tRNA genes and cytoplasmic tRNAs animal mitochondria have only 22 different tRNA types to translate the 20 amino acids [61, 62]. With two exceptions, isoacceptor tRNAs are not available inside animal mitochondria. Leucine and serine each have two mitochondrial tRNAs with different anticodons; the remaining 18 amino acids are each translated by a single tRNA that covers all cognate codons. The best documented mechanism for altering the efficiency of translation is when the changed codon(s) are near to the first-translated end of the gene [54]. The approximately even distribution of synonymous variation among mitochondrial genes in modern humans [34] is most compatible with neutrality [63].

Speculatively, synonymous codon changes could affect gene expression by mechanisms independent of tRNAs. These include changes in mRNA secondary structure and stability and the binding of specific factors, protein or miRNA. Modification of splicing is a candidate and possibly important mechanism by which synonymous codons alter protein structure and function. However, in contrast to most nuclear genes in most animals, animal mitochondrial protein-encoding genes do not have introns. In animal mitochondria there are no alternative spliced forms whose ratios could be modulated by synonymous codons near splice sites.

Kimura’s insight that a preponderance of synonymous substitutions is evidence for neutral evolution [64] now appears to be more universally valid for the mitochondrial than for the nuclear genome. For nuclear genomes one finds a growing number of cases and mechanisms where synonymous codons have phenotypes and are subject to selection. In contrast, for the mitochondrial genomes of animals there is not a single example of any synonymous codon having a phenotype. Furthermore, the known mechanisms that allow synonymous codons to alter the phenotypes of nuclear genes are impossible in mitochondria. Mitochondrial sequences yield straightforward and uncomplicated phylogenetic analysis and species-level identification for reasons beyond those known by those who originally proposed them. Sometimes you get lucky.

## The case of the missing G’s

Codons that end in G are underrepresented by a factor of about ten in animal mitochondria. Previously we interpreted the lack of third position G’s in mitochondrial coding sequences is evidence of a role for extreme purifying selection in determining the COI DNA barcoding gap [20] but we now find this argument flawed. On the one hand there appears to be purifying selection against codons that end in G but this apparent selection is similar in neighboring species. With selection against G-ending codons in all animal species it could not be a source of species-specific adaptive peaks. Further insight into the lack of G in the third position follows from a focused review of the wobble hypothesis in the context of mitochondria.

Francis Crick set forth a set of stereochemical models in which a single tRNA anticodon pairs with multiple codons for the same amino acid. Crick called his idea the “Wobble hypothesis” because it postulated flexible pairing between the 3’ base of the codon with the 5’ base of the anticodon [65]. The Wobble hypothesis is brilliantly insightful, however, details of the pairing scheme have changed with knowledge of extensive post-transcriptional chemical modifications of tRNAs. More than 150 different chemical modifications of RNAs are now characterized [66]; the greatest concentration of RNA modification is found on anticodons. Only certain modifications at the U at the 5’ position in the anticodon allow efficient Wobble-pairing with G [67–70]. Wobble G is rare in animal mitochondrial codons[20] consistent with the fact that Wobble G recognition-specific modifications have not been found in animal mitochondrial tRNAs [71, 72].

Several human pathologies are correlated by Genome Wide Association Studies (GWAS) to SNPs that change a synonymous codon and it is expected that more will be found [73]. Two cautions apply when considering Genome Wide Association Studies linking synonymous codons with human mitopathologies: 1) So far as we are aware there are no inferences of pathologies based on synonymous substitutions in the mitogenome [74], 2) GWAS are subject to artifacts of inference that encourage erroneous confidence [75, 76]. GWAS are hypothesis generators, not proof. Anecdotally, workers in mitochondrial pathologies are well aware of synonymous codons, and so the absence of evidence for any human pathology due to a synonymous codon change in the mitochondrial genome is not due to a lack of looking. In contrast, mutations in mitochondrial tRNA genes are “hotspots” for human pathologies [77]; they lead to large scale insertion of inappropriate amino acids.

## Selectionist arguments

Thomas Kuhn points out that science stalls when different camps that study the same aspects of nature use different vocabularies, cite only within their own discipline and ignore or disparage each other. There appears to be an unfortunate isolation in the literature between camps that advocate mitochondrial selection and those that rely on demographic reasoning. Here we argue that selectionist perspectives are valid in some mitochondrial systems and plausible in others. However, exceptions abound and as a whole we find that a selectionist perspective is not robust enough to account for the animal-kingdom wide facts of the barcode gap.

Even within a single species, different external environments may select for particular alleles of mitochondrial-encoded enzymes [78–81]. In modern humans there are two different amino acid alleles of the mitochondrial-encoded ATPase. This subunit partitions the proton gradient of mitochondria in two ways: it can use the gradient to form the covalent to join inorganic phosphate to ADP in order to make ATP. Alternatively, if the proton gradient runs down without storing energy in the synthesis of ATP heat is immediately released. The allele predisposed to ATP synthesis is more frequent in human populations who inhabit tropical regions. Conversely, the allele biased toward instantaneous heat generation is more frequent in colder regions [82]. The argument for environmentally-driven selection for this allele is logical and inspires interesting experiments [83]. The plausible but unproven possibility of selection for a single allelic case of amino acid substitution is a small pebble in the scale when compared to the evidence for the apparent neutrality of most mitochondrial variation. Most human mitochondrial variation, similar to that of other animal species, consists of synonymous codon changes. However, in principle, the linkage of a single selected amino acid could drive a species-wide sweep of the entire linked mitochondrial genome.

The model of an optimum sequence for each species has two subcategories: a) optimal for the external environment, and b) optimal integration with other genes of the organism [84]. These two mechanisms can work together and various permutations have been suggested with more or less emphasis on selection for external conditions such as environmental temperature or internal compatibility with nuclear genes [80, 85–87]. Compatibility among the thousand or so nuclear genes whose products enter the mitochondria and the 13 gene products coded for by the mitochondrial genome can lead to reproductive isolation and incompatibility [21, 86, 88]. Incompatibility of mitochondrial and nuclear genes can cause reproductive isolation either immediately or via decreased fitness of progeny [87, 89–92]. Mitochondrial introgression in some cases has been proposed to favor the co-introgression of compatible nuclear alleles that form subunits of mitochondrial complexes [93, 94]. This is a potentially important perspective but its generality is unclear.

We see arguments against stepping from specific examples of nuclear-mitochondrial incompatibilities to a general theory of this effect as the major driver of animal speciation: 1) There are many examples of fertile and fit interspecies hybridizations including cases with mitochondrial sequences that diverge by 4% or more [95–102]. 2) Some well-studied cases of mating incompatibility have nothing to do with nuclear-mitochondrial incompatibility. Examples include chromosomal inversion [103, 104] and the activation of endogenous retroviruses-transposons [105]. 3) Nuclear-mitochondrial incompatibility is a subset of physiological incompatibility. Other definitions of species, e.g. behavioral, geographical, ecological, do not require physiological mating incompatibility in any form [11]. COI barcode clustering is more widely a fact than can be accounted for by mating incompatibility in general and nuclear-mitochondrial incompatibility in particular. 5) Finally, there is no example in which mating incompatibility or weakness of inter-species hybrids is attributable to the synonymous codons that constitute the major fraction of barcode gaps.

The average pairwise difference of the COI barcode in modern humans is 0.1%, i.e., about average for the animal kingdom. However, the most extreme differences between individual humans approach 1%. This difference is as great as many distinctions among neighboring species. Modern humans are a single population. Darwin made this point with respect to visible phenotypes and it applies even more strongly when neutral variants are considered:

*Hereafter, we shall be compelled to acknowledge that the only distinction between species and well-marked varieties is, that the latter are known, or believed, to be connected at the present day by intermediate gradations, whereas species were formerly thus connected [106].*

The possibility of preferred combinations of nuclear and mitochondrial alleles within a species is intriguing and there is one example of experimental support. An inbred strain of mouse was shown to have non-optimal physiology when the mitochondrial genome from a different inbred line was crossed in (10 backcrosses to the nuclear line all using the female descendent from the first mitochondrial donor) [107]. This finding has been extrapolated as justification to urge studies of nuclear-mitochondrial compatibility in human three-parent IVF (in vitro fertilization) [108]. On the other hand, human mitochondrial transfer experiments have found no analogous effect [109], arguably, owing to the different genetic structure of our species when compared to inbred mouse strains [110]. The differences in the two mouse mitochondrial genomes at issue include missense in the coding region, tRNA alterations and ori-region changes as well as synonymous codon changes. There are no data to pinpoint which sequences make a difference, in particular no evidence for a phenotype of synonymous codon changes, which the authors mark as “silent” (Extended Data Table 1 in [107]).

This line of work and controversy adds evidence that in some cases mitochondrial-nuclear incompatibility may interfere with mating or health of offspring. However, the work does not show any effect of synonymous codon changes. No matter which mechanism for speciation is responsible in any specific case, the 0.0%-0.5% accumulation of synonymous variance independent of population size or apparent species age is a biological fact. The variable distance between the most closely related living species presumably reflects differing numbers of extinct intermediate sequences.

## Conditions that favor clonal uniformity are frequent in biology

Bottlenecks, founder effects, lineage sorting, and gene sweeps decrease genetic diversity in a population [111]. The question is how widespread these effects are in the context of defining animal species and if it is possible distinguish them in other than a rhetorical manner. Here we emphasize the overlap—in fact the near congruence—in the conditions that favor each of these mechanisms.

Based on contemporary mitochondrial sequence data alone it is impossible to distinguish an organismal bottleneck from mitochondrial and Y chromosome specific lineage sorting since both mechanisms make the same prediction of a uniform mitochondrial sequence in the past [112].

A positively selected allele has the potential to sweep through a population and by hitchhiking [113, 114] or genetic draft [115] carry the entirety of the linked genome along thereby resetting mitochondrial variation to zero. This scenario requires a single maternal lineage replace all others [113]. It is reasonable to hypothesize that somewhere on the mitochondrial genome there arises a positively selected amino acid substitution leading to the replacement of the entire linked genome in the entire population. One should not mince words about what a mitochondrial genome sweep requires: the entire population’s mitochondrial genome must re-originate from a single mother.

These three pathways toward sequence uniformity should not be thought of as enemies because they converge in both cause and effect. Lineage sorting is most efficient when the population is small, when the number of different mitochondrial genotypes is small, and when the population is either stable or shrinking [116]. An extreme diminishment of population size followed by population expansion is the definition of a bottleneck. Lineage sorting is diminished during periods of population growth and does not occur at all during exponential growth when all neutral lineages leave progeny [117, 118]. The same conditions that favor lineage sorting also favor gene sweeps, which in the context of a totally-linked genome means one mitochondrial genome. The concept of “gene sweeps” emphasizes positive selection whereas “lineage sorting” emphasizes neutral events. Bottlenecks are extreme forms of the same conditions.

Bottlenecks followed by expansion are the dominant mechanism for evolution in the microbial majority of life and it might seem odd to think animals should be exceptional [119]. Ever since Koch, microbiologists have streaked out their bacteria to begin experiments with pure, i.e., clonal cultures [120]. The first experiments showing evolution of new mutants from clonal starting populations were the classical cases of proof that bacterial genes follow the patterns expected from random mutation that grow indistinguishably from sibs when unselected ([117, 121–123]). Clonal outgrowth and replacement of the inoculating population was inferred from the earliest chemostat [124] as well as later serial transfer experiments [125]. Epidemiology shows that repeated bottlenecks play dominant roles in the natural evolution of microbial pathogens including protists, bacteria and viruses [126–132]. A visually impressive demonstration of successive clonal selection and population outgrowth is seen in time lapse studies of bacteria serially mutating to new heights of antibiotic resistance [133]. On the host side of the equation, the clonal selection theory of immune system development was controversial when first proposed but its logic and experimental support proved compelling [134, 135].

Mayr made a specific proposal for the role of extreme bottlenecks in speciation that followed from a founder effect (originally 1942, here quoted from a reprise based on interviews in 2004 [136]):

*The reduced variability of small populations is not always due to accidental gene loss, but sometimes to the fact that the entire population was started by a single pair or by a single fertilized female. These “founders” of the population carried with them only a very small proportion of the variability of the parent population. This “founder” principle sometimes explains even the uniformity of rather large populations, particularly if they are well isolated and near the borders of the range of the species.*

Eldredge and Gould used this idea of allopatric speciation in small isolated populations that then rapidly expanded to account for the abrupt transitions seen in the broad range of the fossil record [137].

Models of allopatric or peripatetic speciation invoke a bottleneck with an additional feature: What emerges from the bottleneck looks or acts differently, i.e., it is a bona fide new species. It may be more frequent that what emerges from a bottleneck looks and acts like a middling representative of what went in.

If mitochondria are considered “honorary prokaryotes” then the dominant mode in prokaryotes of frequent processes that lead to clonal outgrowth either by selection or random processes [138] are not counterintuitive. A number of different processes could lead to the mitochondrial sequence becoming clonal. Candidate processes include bottlenecks and lineage sorting on three different levels: Within organelles, among organelles in the same cells, among cells in an organism (particularly in the germ line) and among organisms. Not certain is whether different processes have led to a similar result throughout the animal kingdom or if a single process operates throughout. Occam’s razor, the principle of parsimony, suggests that a single explanation should be considered.

Purifying selection in linked genomes slows but does not stop the accumulation of neutral variation [139]. Drift and lineage sorting during population stasis or shrinkage decrease variation. The efficiency of decrease depends on the number of haplotypes in the population, as well as the numbers and distributions of female offspring among parents with different haplotypes [10]. A key prediction of naïve neutral theory that does not hold up against extensive barcode data from across the animal kingdom is that larger populations or older species should harbor more neutral variation [20, 140, 141]. The key incompatibility of naïve neutral theory with biological fact is that the theory considers populations at equilibrium in the sense that the population be at stable numbers for approximately as many generations as the mutation rate per generation. The evolution of modern humans offers a specific solution to the animal-kingdom-wide dilemma of missing neutral mutations.

## Modern humans

More approaches have been brought to bear on the emergence and outgrowth of Homo sapiens sapiens (i.e., modern humans) than any other species including full genome sequence analysis of thousands of individuals and tens of thousands of mitochondria, paleontology, anthropology, history and linguistics [61, 142–144]. The congruence of these fields supports the view that modern human mitochondria and Y chromosome originated from conditions that imposed a single sequence on these genetic elements between 100,000 and 200,000 years ago [145–147]. Contemporary sequence data cannot tell whether mitochondrial and Y chromosomes clonality occurred at the same time, i.e., consistent with the extreme bottleneck of a founding pair, or via sorting within a founding population of thousands that was stable for tens of thousands of years [116]. As Kuhn points out unresolvable arguments tend toward rhetoric.

## Summary and conclusion

Science greedily seizes simplicity among complexities. Speciation occurs via alternative pathways distinct in terms of the number of genes involved and the abruptness of transitions [148]. Nuclear variance in modern humans varies by loci in part due to unequal selection [149] and the linkage of neutral sites to those that undergo differential selection. Complexity is the norm when dealing with variance of the nuclear ensemble [150–154]. It is remarkable that despite the diversity of speciation mechanisms and pathways the mitochondrial sequence variance in almost all extant animal species should be constrained within narrow parameters.

Mostly synonymous and apparently neutral variation in mitochondria within species shows a similar quantitative pattern across the entire animal kingdom. The pattern is that that most—over 90% in the best characterized groups—of the approximately five million barcode sequences cluster into groups with between 0.0% and 0.5% variance as measured by APD, with an average APD of 0.2%.

Modern humans are a low-average animal species in terms of the APD. The molecular clock as a heuristic marks 1% sequence divergence per million years which is consistent with evidence for a clonal stage of human mitochondria between 100,000-200,000 years ago and the 0.1% APD found in the modern human population [34, 155, 156]. A conjunction of factors could bring about the same result. However, one should not as a first impulse seek a complex and multifaceted explanation for one of the clearest, most data rich and general facts in all of evolution. The simple hypothesis is that the same explanation offered for the sequence variation found among modern humans applies equally to the modern populations of essentially all other animal species. Namely that the extant population, no matter what its current size or similarity to fossils of any age, has expanded from mitochondrial uniformity within the past 200,000 years.

Nonhuman animals, as well as bacteria and yeast, are often considered “model systems” whose results can be extrapolated to humans. The direction of inference is reversible. Fossil evidence for mammalian evolution in Africa implies that most species started with small founding populations and later expanded [157] and sequence analysis has been interpreted to suggest that the last ice age created widespread conditions for a subsequent expansion [158]. The characteristics of contemporary mitochondrial variance may represent a rare snapshot of animal life evolving during a special period. Alternatively, the similarity in variance within species could be a sign or a consequence of coevolution [159].

Mitochondria drive many important processes of life [160–162]. There is irony but also grandeur in this view that, precisely because they have no phenotype, synonymous codon variations in mitochondria reveal the structure of species and the mechanism of speciation. This vista of evolution is best seen from the passenger seat.

## Acknowledgements

Bruce Levin suggested the phrase “honorary prokaryote” in reference to mitochondria. Others have used the phrase in reference yeast or to phage introns. Thanks to Glen Bjork, Manny Goldman, Ken Zahn for discussions on tRNAs, Jesse H. Ausubel, Frank Stahl for encouragement and comments and the Alfred P. Sloan Foundation and Monmouth University/Rockefeller University Marine Science and Policy Initiative for support.

